# ONT-LibQC: rapid quality control of chromatin assay libraries using shallow nanopore sequencing

**DOI:** 10.64898/2026.10.19.760194

**Authors:** Hongjiang Liu, Jongmin J. Kim

## Abstract

A next-generation sequencing (NGS) library for genomic assays can have an acceptable DNA concentration and size distribution yet lack the biological signal of interest. For chromatin assays, such as CUT&RUN and Micro-C, such problems often become apparent only after deep Illumina sequencing, delaying troubleshooting and protocol optimization. Here, we present ONT-LibQC, a workflow that uses a few thousand Oxford Nanopore Technologies (ONT) sequencing reads to examine finished libraries before deep sequencing. It reconstructs library molecules, identifies adapters and indexes, and assesses fragment sizes and the enrichment of biological signal. For CUT&RUN libraries, shallow, approximately 3,000 ONT reads were sufficient to show enrichment of promoter-associated H3K4me3, gene-body H3K36me3 and broad H3K27me3 profiles, consistent with profiles from Illumina deep sequencing. For Micro-C libraries, cis-contact fractions from shallow ONT sequencing closely matched Illumina results. Furthermore, full-length ONT reads revealed internal ligation structure and enabled estimation of the lengths of MNase digested DNA fragments. ONT profiles were reproducible across four technical replicate sequencing runs of each of two H3K4me3 libraries. Downsampling analysis further showed that key quality control (QC) features remained informative at lower read counts as low as 1,000 reads. Together, these results show that shallow ONT sequencing can provide rapid feedback on library quality, helping researchers identify potential problems before committing to deeper sequencing. ONT-LibQC is available through a web portal, a local graphical interface and a command-line workflow.

## Introduction

Recent advances in chromatin-profiling have made it possible to study gene regulation in rare cells (1–3), primary tissues(4, 5), and archived samples(6–8). Applying these assays to new sample types often requires optimization of cell or nuclei preparation(9, 10), fixation(11), digestion(12, 13). A major bottleneck is that an experiment may appear to have produced a suitable sequencing library, yet its success becomes clear only after sequencing and analysis, which can take several weeks. When libraries are submitted to a shared core facility or commercial sequencing service provider, this delay can substantially slow the cycle of troubleshooting and protocol refinement.

Library quality is typically assessed in two stages. DNA quantification and electrophoresis assess whether a finished library contains sufficient material with an appropriate size distribution. Sequencing then reveals whether the library contains the expected biological signal. However, the first stage results cannot reliably answer the second question: a plausible library quantity and profile can be obtained without enriching for the desired signal(14, 15). An intermediate assessment of finished libraries could help researchers identify problems before committing to deep Illumina sequencing.

Available approaches address parts of this gap. For CUT&RUN and related chromatin-profiling assays, qPCR can test selected genomic regions, but it requires information on suitable target and control loci in advance and provides limited information about the broader distribution of signal(16). For chromosome conformation capture assays, such as Micro-C, the finished library size profile provides useful quality control (QC) information about the lengths of ligated DNA fragments, which can reveal differences in digestion and ligation efficiency. However, whether biologically meaningful proximity contact was captured cannot be determined without sequencing steps. Low-depth sequencing(15) using a small-scale Illumina sequencing platform such as the MiSeq could be a solution, but their cost-to-sequence output is low, and the machine is not readily available for most labs. Alternatively, development of modified techniques, such as R2C2 and associated computational method, PLNK, allows accurate sequencing of Illumina libraries using the relatively accessible Oxford Nanopore Technologies (ONT) platform(17). However, the R2C2-PLNK approach requires several additional library-preparation steps, making it less practical for routine QC of a small number of libraries.

Advancement of cost-effective and rapid commercial ONT sequencing services worldwide offers a sweet spot for a rapid, intermediate QC opportunity(18, 19). Some services can return a few thousand full-length reads from a small library aliquot within one or two business days. By employing these accessible sequencing infrastructures, we aimed to develop a quality-control workflow for shallow sequencing of finished chromatin-profiling and Micro-C libraries, with outputs organized around practical experimental questions.

Here we present ONT-LibQC and evaluate which measurements remain informative when as few as hundreds to a few thousand ONT reads are available. We show that focal enrichment patterns across metagenes clearly distinguish high-quality from failed CUT&RUN libraries and reflect patterns obtained by deep sequencing. In Micro-C, shallow ONT sequencing similarly recapitulates key ligation statistics measured by deep sequencing. Replicate sampling demonstrates reproducible QC metrics, and downsampling further shows that as low as 1,000 reads are sufficient to assess key library features. We provide ONT-LibQC as an accessible QC platform with both a web-based graphical interface and a downloadable implementation.

## Results

### Motivation and design of ONT-LibQC: a cost-effective method for next-day evaluation of chromatin profiling assays

The primary goal of ONT-LibQC is to provide rapid assessment of whether chromatin profiling libraries contain the expected experimental features before deep sequencing. The conventional workflow of chromatin profiling experiments is as follows (Fig. 1A). The researcher confirms the concentration and size distribution of the finished libraries using sensitive, automated electrophoresis systems, such as TapeStation and Bioanalyzer. If acceptable, libraries are submitted to sequencing facilities for deep short-read sequencing, most commonly by Illumina platforms. Many laboratories rely on core facilities or commercial sequencing services, which can take 2-4 weeks to return sequencing results. Consequently, potential problems with the experiments may only become apparent several weeks after library preparation. ONT-LibQC provides a low-cost, rapid checkpoint between library preparation and deep sequencing, by adding shallow ONT sequencing of an aliquot of the finished library (Fig. 1A).

**Figure 1.**
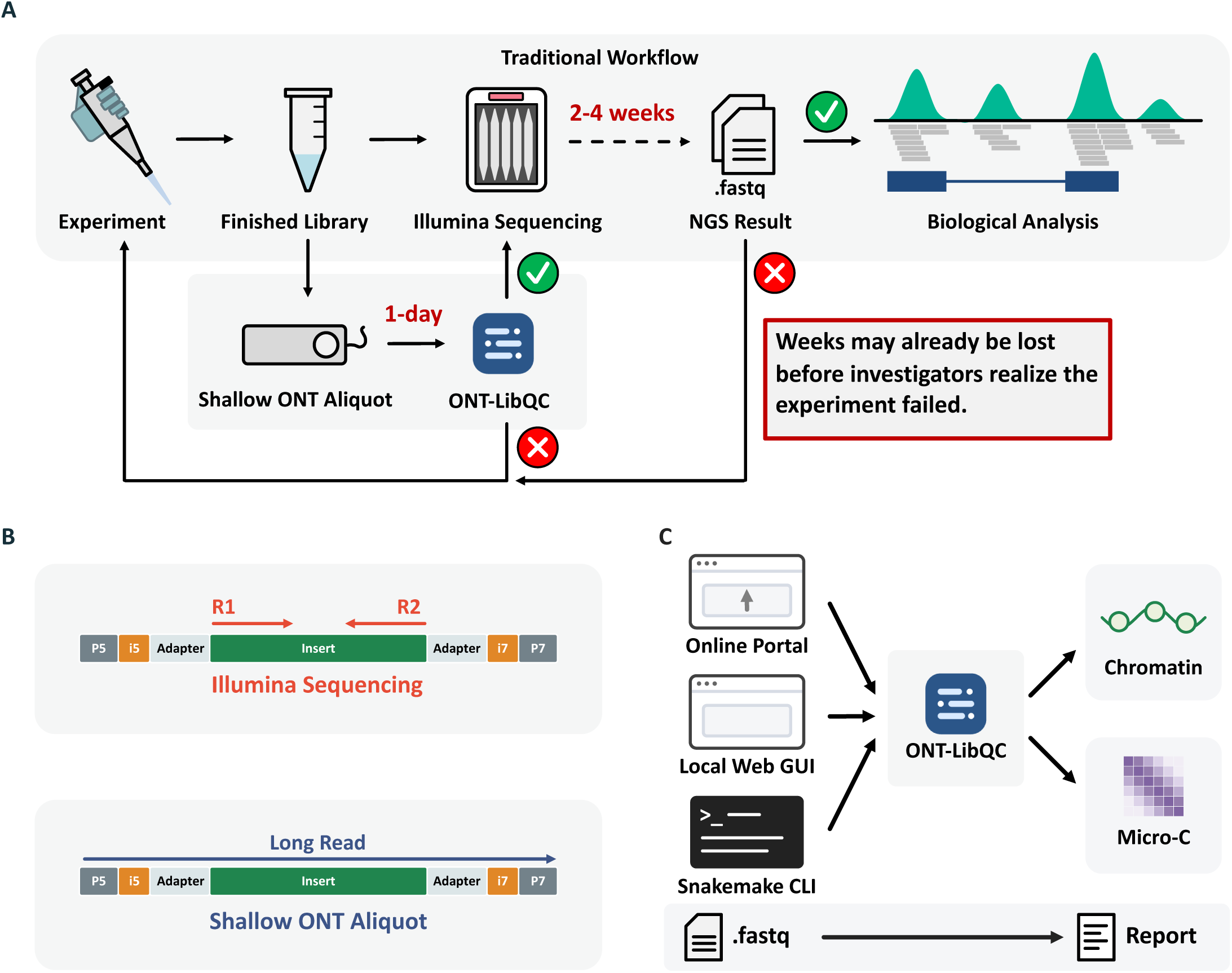
An intermediate QC checkpoint for finished sequencing libraries. **A,** Shallow ONT sequencing allows a finished-library aliquot to be examined before deeper Illumina sequencing. **B,** Paired-end Illumina sequencing reads two ends of a library molecule, whereas a spanning ONT read can reveal adapters, indexes and the intervening insert. **C,** A shared analysis engine provides structural and assay-specific outputs through hosted, local and command-line interfaces.

Commercial ONT amplicon sequencing services can use ONT native barcoding(20), which has two advantages for pre-deep sequencing QC. First, ONT library preparation does not require an additional PCR amplification step, avoiding the introduction of additional PCR bias. Second, full-length molecules are sequenced, allowing adapter and index information to be recovered together with the internal target sequences (Fig. 1B).

We built a computational pipeline, ONT-LibQC, that provides an integrated analysis framework for extracting multiple library-quality metrics from shallow ONT sequencing. The ONT-LibQC analysis platform can be accessed through the hosted portal, a self-hosted browser GUI, or the Snakemake command line (Fig. 1C)(21). Using ∼3,000 ONT reads, users can rapidly obtain general information about library structure, including adapter type, index pair sequences, adapter dimer-like proportions, and fragment length distributions. For chromatin feature enrichment assays, such as ChIP-seq, CUT&RUN, and ATAC-seq, ONT-LibQC provides enrichment profiles over standard gene models or user-provided regions in BED format. For chromatin conformation assays, such as Micro-C, ONT-LibQC provides key ligation statistics.

### Reconstruction of electrophoretic fragment profiles from raw sequence reads

To assess how faithfully ONT sequencing represents the fragment-size distribution of the original library despite the additional library preparation step, we reconstructed electropherogram-like profiles from raw read lengths (Fig. 2). Because electrophoretic fluorescence signal reflects DNA mass rather than molecule count, the contribution of each read to the reconstructed library signal was weighted by its fragment length (see Methods, Reconstruction of native electrophoresis). Across five libraries representing Micro-C and CUT&RUN experiments using *Drosophila* and human cells, ONT-LibQC reconstructed profiles closely recapitulated the corresponding TapeStation electropherograms (Fig. 2). Although larger fragments were underrepresented in the ONT-LibQC profiles, the 300-500 bp range showed close agreement between ONT-LibQC and TapeStation profiles (Fig. 2C and 2D). This size range contains the predominant sequenceable fragments in many Illumina chromatin-profiling libraries.

**Figure 2.**
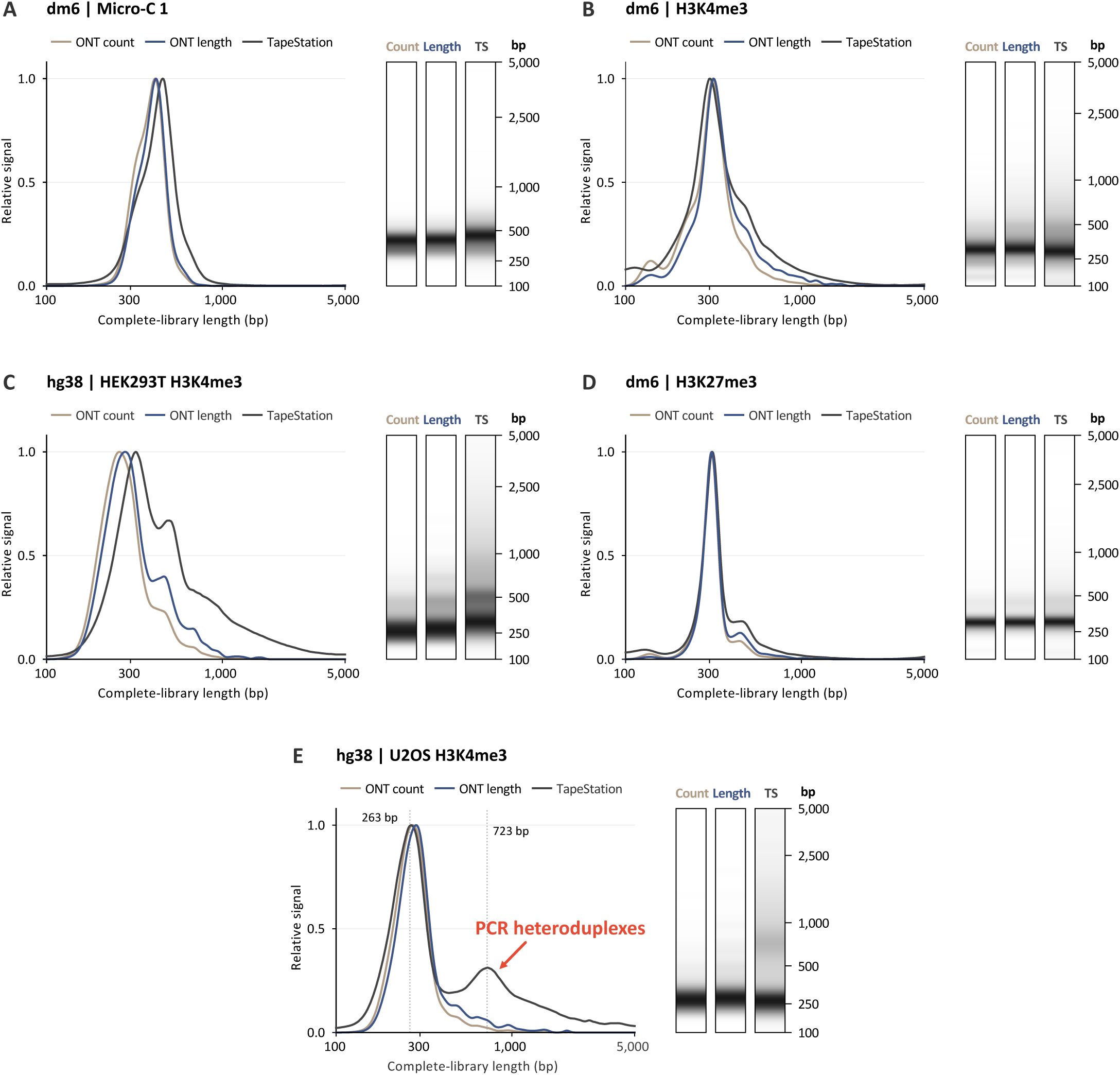
Comparison of ONT-reconstructed size profiles with TapeStation measurements. **A-E,** Count- and length-weighted ONT profiles compared with TapeStation traces for (A) dm6 Micro-C, (B) dm6 H3K4me3, (C) HEK293T H3K4me3, (D) dm6 H3K27me3, and (E) U2OS H3K4me3 libraries. Complete-library lengths were calculated by adding 136 bp of adapters and indexes to the reconstructed insert lengths. Each unit contributed equally to count-weighted profiles or in proportion to its length to approximate DNA mass in length-weighted profiles. Curves were independently peak-normalized within the 100-5,000 bp range. Adjacent gel-like lanes show the same profiles using a shared, ladder-calibrated migration coordinate. Count, count-weighted ONT; Length, length-weighted ONT; TS, TapeStation. In E, the secondary TapeStation peak at an apparent size of 723 bp is consistent with PCR heteroduplexes (bubble products) and has no corresponding peak in the ONT reconstructed profile. Numerical source data are provided in Supplementary Table S2.

A key advantage of the reconstructed ONT-LibQC profile is that the sequences underlying any size range can be directly examined. This is particularly useful for investigating unexpected peaks, such as adapter dimers and PCR heteroduplexes. Adapter dimers are typically approximately 120-150 bp molecules that contain sequencing adapters but lack genomic inserts. Because they can bind and cluster on Illumina flow cells, adapter dimers consume sequencing capacity(22). ONT-LibQC therefore automatically identifies adapter-dimer molecules and reports their abundance. In addition, PCR heteroduplexes, sometimes referred to as “PCR bubbles,” can form when DNA strands containing different insert sequences anneal through shared adapter and index sequences(23). These heteroduplexes migrate more slowly during electrophoresis and can produce artifactual peaks, particularly after excessive PCR cycling or primer depletion(24). The second peak at apparent 723 bp observed by TapeStation has no corresponding peak in the ONT-derived length profile (Fig. 2E), consistent with PCR heteroduplexes rather than long library fragments. Because ONT directly measures the sequence length of each individual molecule, a peak seen by electrophoresis but absent from the sequence-based reconstruction can be flagged as a possible migration artifact.

### ONT-LibQC assesses key library quality features using average signal profiles across metagenes or user-defined regions

Current cost-effective ONT-sequencing services offer 3,000-5,000 reads, which is not sufficient to reliably evaluate enrichment at individual loci. We therefore asked whether averaging signal across many genes or user-defined genomic features, such as reported binding sites, could reveal the expected enrichment pattern. We first tested this using H3K4me3 CUT&RUN datasets, which are expected to show enrichment near transcription start sites (TSSs) (Fig. 3A - 3C, H3K4me3)(25). We analyzed both *Drosophila* and human libraries to determine whether aggregate enrichment could be detected across species with genome sizes differing by approximately 20-fold. When 3,000-5,000 ONT reads were plotted as average metagene profiles, successful libraries clearly showed TSS enrichment, whereas this pattern was not observed in failed libraries for either *Drosophila* or human H3K4me3 datasets (Fig. 3D and 3E). Average profiles generated from deep Illumina sequencing closely matched those obtained from shallow ONT sequencing (Fig. 3D and 3E, ONT-Illumina correlation *Drosophila*: 0.945 and human: 0.922), indicating that the expected enrichment pattern could be recovered from shallow ONT sequencing. In addition, shallow ONT sequencing of an H3K36me3 CUT&RUN library recovered the expected 3’ gene-body enrichment toward the transcription end site (TES)(25), with ONT-Illumina correlation of 0.95 (Fig. 3F). Together, these results indicate that shallow ONT sequencing can reveal whether a finished library contains the expected biological signal.

**Figure 3.**
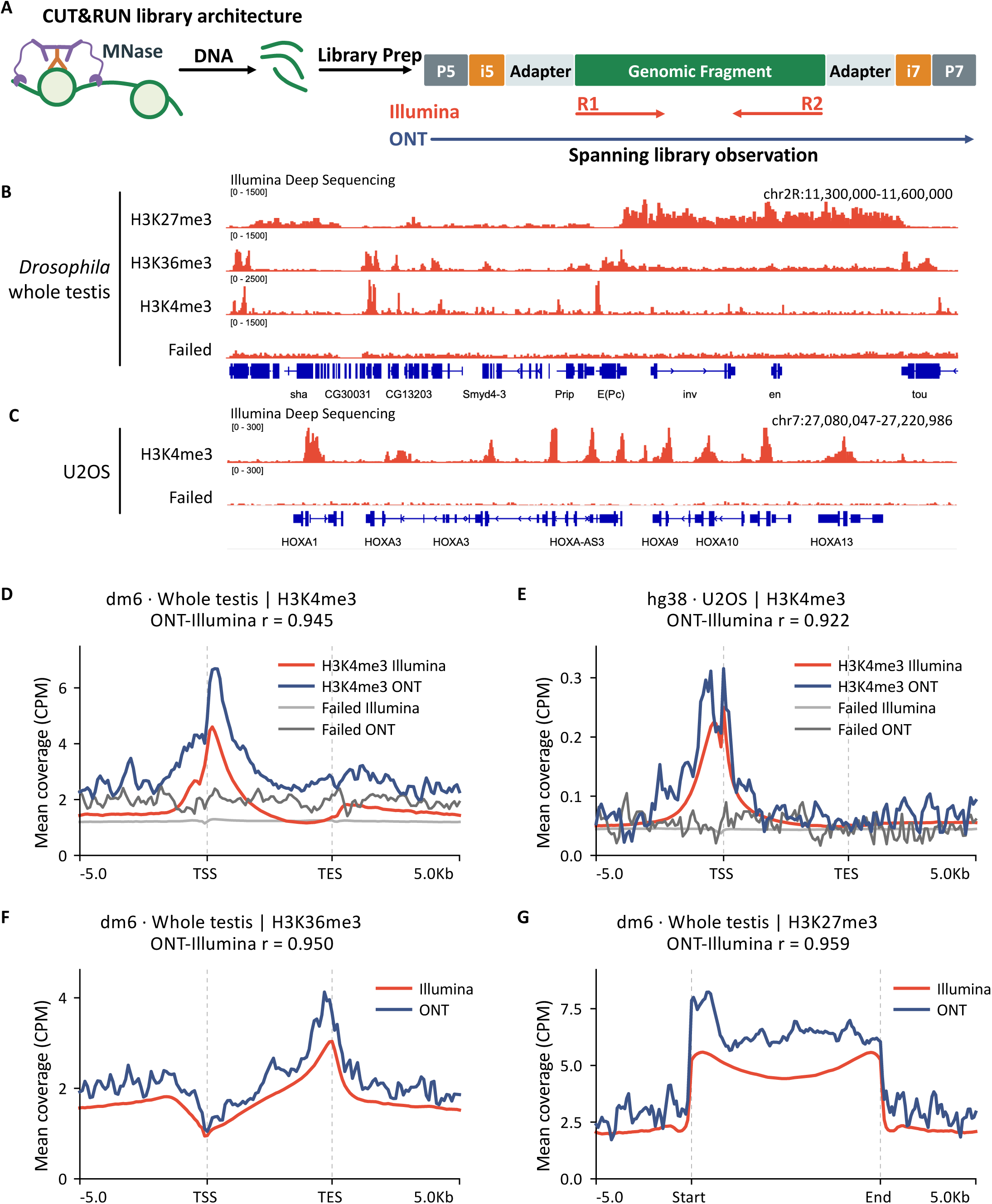
Library structure and target-associated QC of CUT&RUN libraries. **A**, CUT&RUN library architecture and sequencing strategy. Antibody-guided MNase releases target-associated DNA, which is converted into indexed libraries. Illumina sequencing reads the two ends, whereas spanning ONT read can recover the complete library unit. The schematic is not to scale. **B,C,** Representative genome-browser tracks for the indicated CUT&RUN targets and failed libraries from whole *Drosophila* testes (dm6) and U2OS cells (hg38), respectively. **D,E,** CPM-normalized gene profiles of H3K4me3 and failed libraries from whole Drosophila testes (dm6) and U2OS (hg38), respectively. **F,** H3K36me3 gene profiles. **G,** H3K27me3 profiles across custom BED regions corresponding to broad peaks called from the matched Illumina library using MACS3. Illumina references were generated from complete datasets. r denotes whole-profile correlation. Failed libraries represent unsuccessful target-specific preparations, not antibody controls.

Many chromatin regulators localize to gene-distal regions or form broad domains spanning multiple genes, where their enrichment cannot be meaningfully visualized using a standard metagene reference(26). Thus, we introduced a feature in ONT-LibQC that allows users to provide predefined genomic regions in BED file format. To test region-based QC of a broadly enriched histone mark, we analyzed a *Drosophila* H3K27me3 library using broad enrichment regions defined from the matched Illumina dataset. The average ONT profile generated from shallow ONT sequencing showed clear enrichment of H3K27me3 within the defined regions, with signal declining into the surrounding flanks (Fig. 3G). Average profiles generated from shallow ONT and deep Illumina sequencing were highly concordant (Fig. 3G, correlation 0.959), showing that shallow ONT sequencing can recapitulate the aggregate enrichment pattern detected by deep Illumina sequencing. This example illustrates how custom regions extend QC beyond promoter- and gene-centered targets.

We next asked whether genomic concentration of reads, the degree to which sequencing signal is confined to specific genomic regions, could provide a single quantitative metric of library quality. We used a collision observed-to-expected (O/E) ratio based on Morisita’s dispersion index to reflect the nonuniformity of read distribution across the genome(27) (see Methods, quantification of genomic signal concentration). The *Drosophila* H3K4me3, H3K27me3 and H3K36me3 libraries had higher collision O/E ratios than the failed libraries (Extended Data Fig. 1A). However, for human libraries, successful H3K4me3 libraries and failed libraries did not show substantially different collision O/E ratio values despite markedly different promoter profiles (Extended Data Fig. 1B). Thus, concentrated genomic coverage alone did not establish target-specific enrichment and was not sufficient as a universal measure of library quality. Interpreting signal concentration together with assay-relevant profiles provides a more informative assessment of library quality than either measurement alone.

### ONT-LibQC recapitulates contact composition and reveals ligation structure in Micro-C libraries

Chromosome conformation capture approaches, including Hi-C and Micro-C, fragment chromatin and ligate DNA ends that are spatially proximal, allowing chromatin contacts to be inferred from the resulting ligation products (Fig. 4A) (28, 29). Thus, important indicators of library quality include the relative abundance of cis (same chromosome) and trans (different chromosomes) contacts, as well as the fraction of long-range cis contacts that likely capture higher-order chromatin interactions (Fig. 4A). We tested whether shallow ONT sequencing could reproduce the broad contact composition observed in matched deep Illumina sequencing data using four Micro-C datasets from *Drosophila* cells and tissues with different contact profiles. Cis-contact fractions differed by no more than two percentage points between shallow ONT and matched Illumina data (Fig. 4B). Both approaches identified Micro-C 1, 3, and 4 as having high cis contact fractions, whereas Micro-C 2 contained a lower fraction of cis-contacts (Fig. 4B). The fraction of long-range (≥10 kb) cis contacts showed relatively lower concordance between shallow ONT and deep Illumina libraries, although the differences were within 0.7-5.4 percentage points (Fig. 4C). The lower concordance likely reflects greater sampling variability for less abundant long-range cis contacts at shallow sequencing depth.

**Figure 4.**
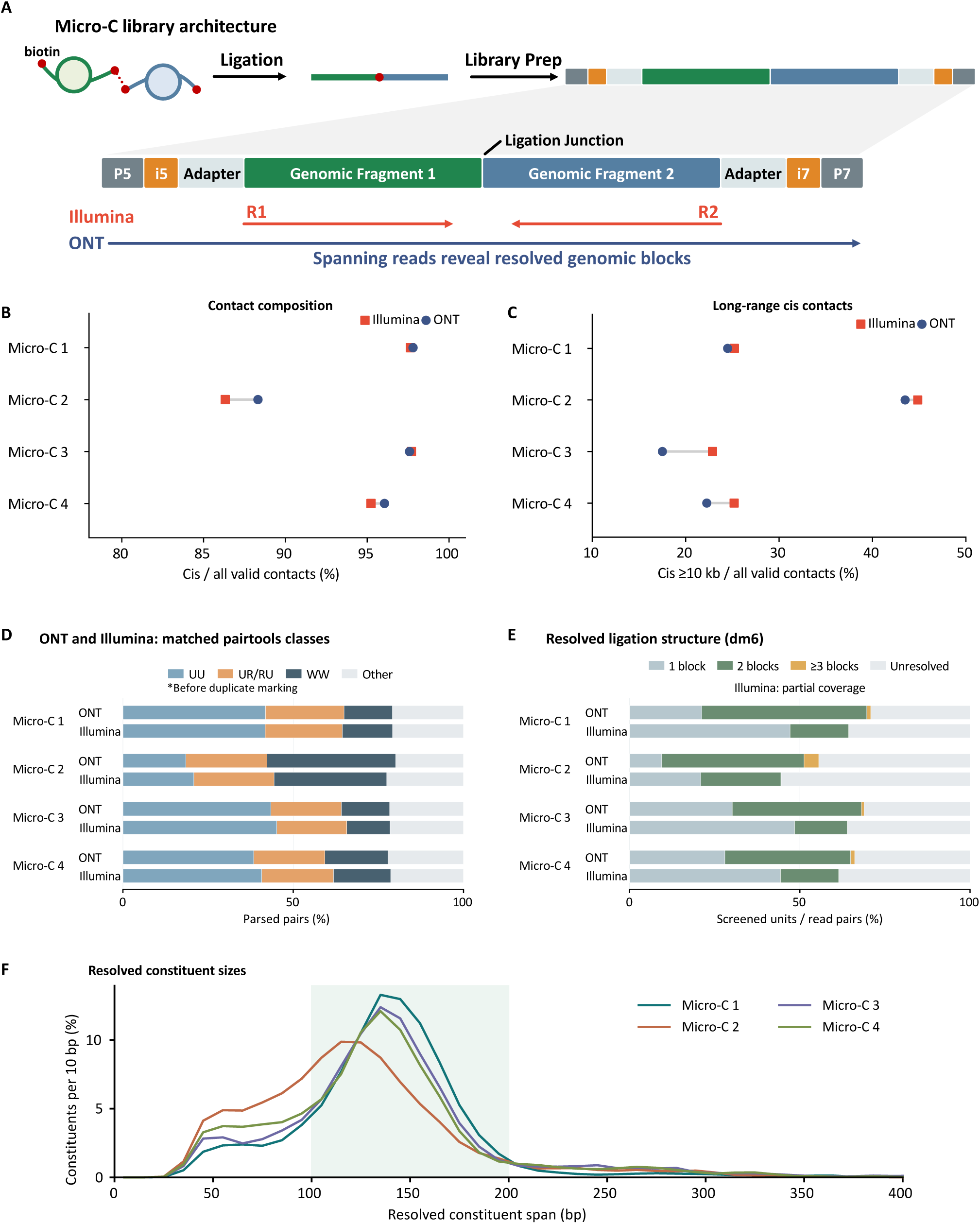
Evaluation of library architecture and complementary QC metrics using *Drosophila* Micro-C data. **A**, Micro-C library architecture, illustrating biotin-labeled fragment ends, proximity ligation, and adapter/index addition. Illumina sequencing reads the ends of ligation products, whereas spanning ONT reads can resolve internal genomic blocks. The schematic is not to scale. **B,C,** Fractions of (B) cis and (C) long-range cis contacts (≥10 kb) from four libraries, calculated using all unique valid contacts under matched filtering criteria. **D,** ONT and Illumina pairtools classifications using a common paired-end analysis protocol without duplicate marking. **E,** Minimum numbers of genomic blocks resolved per reconstructed ONT insert or Illumina read pair. ONT counts use full-length inserts, Illumina counts use paired-end reads. Adapter dimers were excluded from the ONT analysis. **F,** Distributions of constituent reference spans from resolved units containing at least two genomic blocks. Shading indicates 100-200 bp range.

A key advantage of ONT sequencing is that full-length molecule sequencing can reveal the internal structure of ligation products that may not be directly captured by sequencing only the two endpoints using short-read Illumina sequencing(30). We first compared the distribution of pair types reported by pairtools(31), including pairs in which both sides mapped uniquely (UU), pairs with one uniquely mapped side and one side rescued from an ambiguous alignment using the rest of the read (UR/RU), and complex walks, meaning reads with more than two alignments (WW). The proportions estimated from shallow ONT sequencing closely matched those obtained from deep Illumina sequencing, with each pair-type category differing by no more than 4.7 percentage points (Fig. 4D). We next examined the number of distinct genomic blocks represented within individual ONT molecules, which can reflect the complexity of ligation events. Approximately 38-50% of screened molecules contained at least two resolved genomic blocks, consistent with ligation of genomic fragments originating from distinct genomic positions (Fig. 4E, Extended Data Fig. 2A). Two-block molecules predominated, while three-block molecules were detected in all four libraries and four-block molecules in two libraries, representing more complex ligation patterns (Fig. 4E, Extended Data Fig. 2A). These observations provide an additional view of ligation complexity that is not directly captured by conventional endpoint contact summaries.

In addition, full-length sequencing allows inference of the length of resolved fragments within ligation products (Fig. 4F). Most inferred fragment lengths were 100-200 bp (Fig. 4F, shaded area), whereas one library (Micro-C 2) showed a prominent shift toward shorter fragments. Such patterns could guide investigation of digestion conditions. One limitation of ONT-LibQC is that it was not suitable for estimating library complexity due to its extremely low sampling depth(32). Consistent with this, duplicate pairs (DD, pairs flagged as duplicates by pairtools) were uncommon in the shallow ONT datasets (Extended Data Fig. 2B), as the probability of sampling the same library molecule more than once is low at this sequencing depth. However, libraries that pass ONT-LibQC can subsequently be evaluated by additional shallow Illumina sequencing to estimate library complexity, which is often performed before committing to the deeper sequencing required for Micro-C studies(33). Together, contact composition and ligation structure profiles provide complementary measures of Micro-C library quality.

### Enrichment profiles were reproducible across technical replicate sequencing runs

The 3,000-5,000 reads obtained by shallow ONT sequencing represent well below 0.1% of the sequencing depth commonly used for many chromatin profiling experiments(34, 35). We therefore asked whether sampling only ∼3,000 ONT reads could provide reproducible estimates of library characteristics, by performing multiple replicate shallow ONT sequencing of the same libraries. Two human H3K4me3 CUT&RUN libraries were each independently sequenced in four technical replicates and compared with their corresponding deep Illumina sequencing datasets. Promoter enrichment profiles were reproducibly observed across all four ONT datasets (Fig. 5A and 5B), with ONT-Illumina profile correlations ranging from 0.874 to 0.948 (Fig. 5C-5D). Length-weighted fragment size profiles were also similar across technical replicates, with modest variation in peak position and the representation of longer fragments (Fig. 5E and 5F). These results indicate that key library-quality features can be reproducibly assessed from ∼3,000 shallow ONT reads.

**Figure 5.**
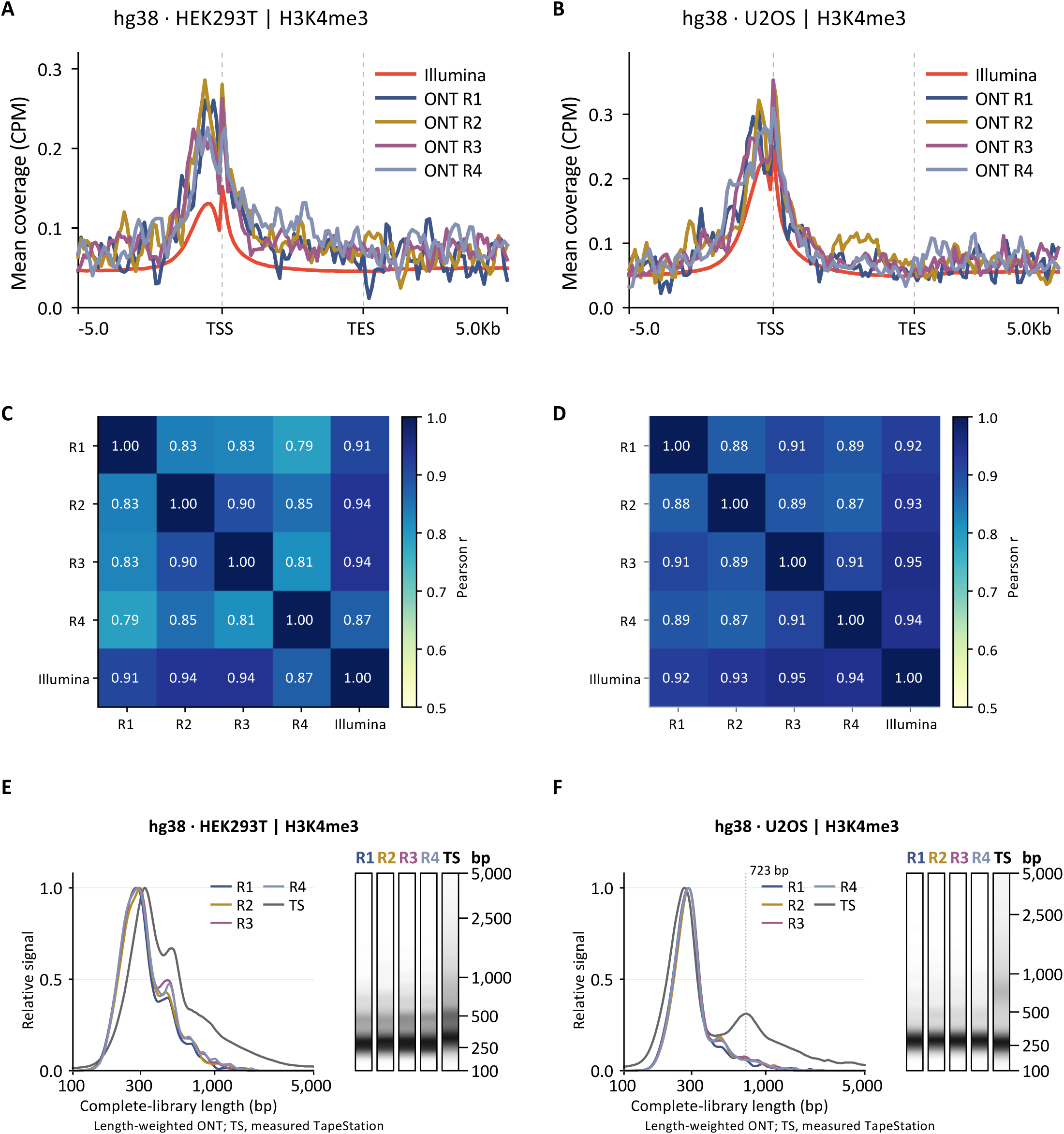
Reproducibility of H3K4me3 CUT&RUN profiles across independent ONT sequencing runs. **A,B,** Four independent ONT sequencing runs of the same (A) HEK293T and (B) U2OS libraries (hg38), each compared with the corresponding Illumina reference using CPM-normalized gene profiles. **C,D,** Pearson correlations among the four ONT profiles and their corresponding Illumina reference profiles. **E,F,** Length-weighted ONT size profiles and corresponding gel-like lanes for the (E) HEK293T and (F) U2OS libraries. Each series includes a single measured TapeStation reference (TS). The 723-bp annotation in (F) indicates the apparent size of the secondary TapeStation peak, which lacks a corresponding peak in the ONT size profile and is consistent with PCR heteroduplexes (bubble products). Curves and lanes were independently peak-normalized within the 100-5,000bp range; right-hand ticks indicate fragment sizes on the ladder-calibrated migration coordinate. The ONT datasets were generated from separate sequencing batches of the same finished libraries and do not represent biological replicates.

### Downsampling analyses show that ∼1,000 reads are sufficient to assess key library features

To determine the minimum read depth required for reliable library evaluation, we computationally and experimentally downsampled a human H3K4me3 CUT&RUN library and performed the same analyses used for our ∼3,000 read ONT datasets (Fig. 6A). For computational downsampling, we generated 50 replicate subsampling series containing nested datasets of 250, 500, 1,000, and 2,000 reads, such that each larger subsample contained all reads from the corresponding smaller subsample. Across 50 independent subsamples of 1,000 ONT reads, the median correlation between each subsampled enrichment profile and the Illumina profile was approximately 0.8 (Fig. 6B). At lower read depths, the median correlations decreased to 0.69 at 500 reads and 0.57 at 250 reads (Fig. 6B). We also tested downsampling experimentally by mixing the H3K4me3 library with increasing amounts of an unrelated library, thereby reducing the number of H3K4me3 reads recovered by ONT sequencing. This experimental dilution generated datasets containing 506, 1,081, and 1,805 H3K4me3 ONT reads, which showed correlations of 0.79, 0.87, and 0.90, respectively, with the corresponding Illumina enrichment profile (Fig. 6C). The correlation between the original undiluted library with the Illumina profile was 0.92 (Fig. 6D). Together, these results suggest that the TSS enrichment pattern remains reproducible at approximately 1,000 reads, whereas lower read depths progressively reduce concordance with the Illumina profile.

**Figure 6.**
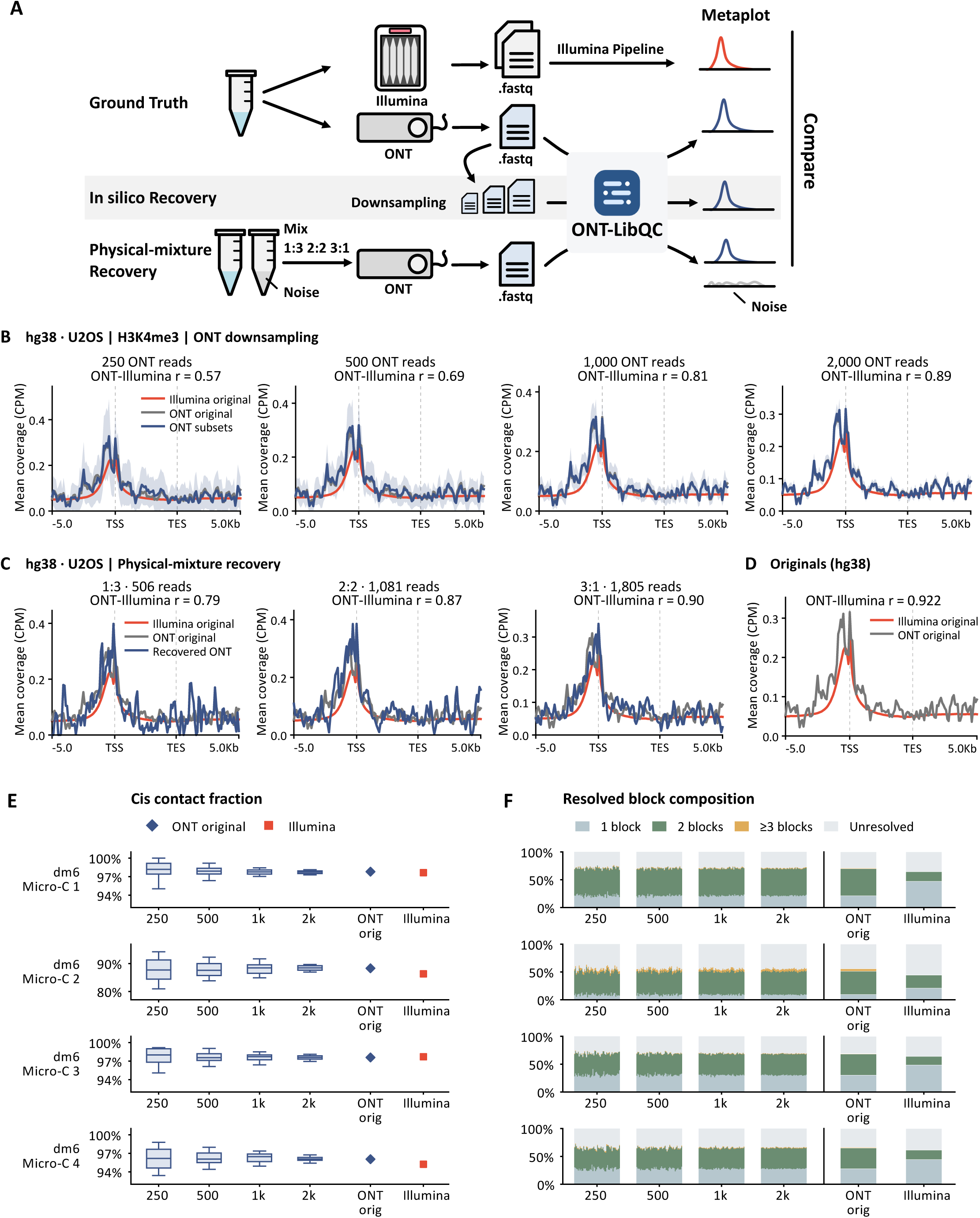
Computational downsampling and experimental library mixing analyses assess the robustness of ONT-LibQC. **A,** Schematic illustrating the comparison of ONT computational downsampling and experimental library mixing results with Illumina reference profiles. **B,** H3K4me3 CPM-normalized profiles (hg38), showing mean ± SD across 50 nested random draws. r indicates the median correlation of individual draws with the Illumina reference. **C,** H3K4me3 profiles from experimental library mixtures, labeled by the number of assigned raw-read counts. **D,** Profiles from the original, unmixed libraries. **E,** Micro-C cis-contact fractions (dm6), showing medians, interquartile ranges, and 5th-95th percentile whiskers, with separate points for the original ONT and Illumina datasets. **F,** Full-insert ONT block counts and partial-coverage Illumina references; each thin bar represents one ONT random draw. Corresponding pairtools processing classes are shown in Extended Data Figure 2.

For Micro-C, we computationally downsampled ONT datasets and evaluated three QC measurements: cis-contact fraction, the number of genomic blocks represented within individual ligation products, and the distribution of pair types reported by pairtools. For cis-contact fraction, the median values across 50 downsampling replicates remained within approximately ±3 percentage points of the value measured from the corresponding Illumina deep sequencing library at all tested depths of 250, 500, 1,000, and 2,000 reads (Fig. 6E). At 1,000 reads, the 5th-95th percentile interval also became narrow, spanning less than 5 percentage points for all but one library (Fig. 6E). Similarly, the distribution of genomic block number and pair types showed relatively little variation among the 50 subsamples once the read depth reached approximately 1,000 reads (Fig. 6F and Extended Data Fig. 2C).

Together, these analyses indicate that the key CUT&RUN and Micro-C QC features examined here can be reproducibly assessed with approximately 1,000 reads. This is substantially below the ∼3,000-read minimum typically provided by many commercial ONT sequencing services, suggesting that the sequencing depth used for ONT-LibQC provides sufficient sampling for these QC measurements.

### ONT-LibQC provides accessible interfaces and integrated reports for library quality assessment

To make shallow ONT-based library QC accessible to users with different levels of computational expertise, we implemented ONT-LibQC as a hosted web application, a self-hosted web interface, and a Snakemake command-line workflow (Fig. 1C). In the hosted interface, users upload FASTQ files and specify the reference assembly and assay type for each library. Each submitted file is analyzed independently and generates a dedicated task page, with an optional email notification upon completion (Fig. 1C). The analysis also includes additional QC features, including a screen for potential spike-in DNA (Extended Data Fig. 3) and mitochondrial fractions (Extended Data Fig. 4). The resulting HTML report, figures, numerical tables, and analysis metadata are packaged together in a single output directory. The self-hosted web interface and Snakemake workflow use the same underlying analysis pipeline, allowing identical analyses to be performed locally or incorporated into larger computational workflows.

Together, ONT-LibQC converts a few thousand shallow ONT reads into an integrated assessment of library structure and assay-specific signal. By providing these measurements before committing libraries to deeper sequencing, ONT-LibQC creates a quick intermediate QC checkpoint at which potential experimental problems can be identified and libraries can be prioritized for deeper sequencing or further troubleshooting.

## Discussion

ONT-LibQC provides a cost-effective and rapid checkpoint between library preparation and deep sequencing. It examines several aspects of a finished library together: its molecular size and structure, enrichment of assay specific signals, and fragment composition for 3C type assays. By integrating these assessments, ONT-LibQC gives users an early, comprehensive view of library quality, well before deep sequencing results become available.

Sequence-based reconstruction of electropherograms provides a secondary confirmation of library size distribution, while also providing detailed information derived directly from the sequence data. Weighting fragments by length improved agreement between ONT-reconstructed and TapeStation profiles, because TapeStation fluorescence reflects DNA mass rather than fragment counts. It therefore provides a suitable basis for visualizing sequence-derived library-size profiles alongside native electropherograms. ONT-based reconstruction retains the sequence information underlying each molecule. While adapter dimers and PCR heteroduplexes are typically inferred from electrophoretic migration patterns alone, ONT-LibQC offers a sequence-based view of the same library. Adapter dimers can be identified and quantified directly from read content. In contrast, the absence of an ONT length peak matching an electrophoretic peak is only indirect evidence: it is consistent with a heteroduplex artifact but does not prove one.

Average chromatin profiles were informative despite shallow coverage at individual loci. H3K4me3, H3K27me3 and H3K36me3 illustrate the value of selecting gene or custom-region coordinates appropriate to the target. Given the nature of shallow sampling, ONT-LibQC is expected to generate clearer signals when enrichment is concentrated or shows a strong feature-aligned pattern, and when the genome size is smaller. In contrast, extremely broad targets with relatively diffuse genomic enrichment, such as H2AK119ub1 or H3K27me2(36, 37), may be less suitable for assessment by shallow sequencing. Thus, the utility of shallow sequencing is expected to depend on both the concentration of the target signal and the size of the genomic search space.

Micro-C illustrates both the strengths and limitations of shallow ONT sequencing-based QC. Because duplicate pairs are rarely sampled at this depth, ONT-LibQC cannot estimate library complexity, and a low duplicate fraction should not be taken to indicate low duplication in the deeply sequenced library(38). Nevertheless, shallow ONT closely reproduced the overall cis-contact composition and provided complementary information on internal ligation structure and constituent-fragment sizes, supporting its use for assessing broad contact composition and recoverable library structure.

Spike-in representation provides another useful dimension of library QC. Robust normalization of chromatin profiling experiments requires appropriate spike-in/sample proportions and sufficient spike-in coverage. Variation in the initial spike-in/sample chromatin ratio or too little spike-in DNA can distort interpretation if it is not accounted for. Sequencing unenriched input controls has been suggested to assess this ratio(39). ONT-LibQC can report the ratio of qualifying cross-species spike-in inserts to host inserts, which can help identify insufficient spike-in read fraction or excessive sequencing reads to spike-in material. However, the spike-in fraction observed after enrichment can also be influenced by assay-specific or biological enrichment and therefore should not be interpreted as a direct measurement of the initial spike-in/sample mixing ratio.

Several practical strategies could further reduce the sample and sequencing requirements of ONT-LibQC. When library material is limited, researchers could further PCR amplify a small aliquot of the libraries for ONT-LibQC, preserving most of the original library for Illumina sequencing. Additional PCR cycles may, however, introduce amplification bias(40). In addition, libraries with distinct index pairs could be pooled for shallow ONT sequencing and demultiplexed by ONT-LibQC, potentially reducing the sequencing cost per library, provided that sufficient usable reads are recovered for each library.

The present comparisons evaluate technical repeatability and sampling stability within the tested libraries. For future development, the reconstruction workflow can be adapted to accommodate predefined library architectures, including designs with unique molecular identifiers (UMIs) following index sequences. This flexibility could extend ONT-LibQC to other indexed short-read libraries, including single-cell RNA sequencing, spatial transcriptomics, and CLIP-seq, with assay-specific validation of reconstruction accuracy, QC metrics and sequencing-depth requirements. For RNA-related libraries, the metaplot framework could be extended by scaling the 5′ UTR, coding sequence and 3′ UTR separately to generate transcript structure-level profile. Thus, ONT-LibQC has potential to be further developed into an adaptable checkpoint for assessing diverse library structure and assay-associated signal beyond the initial presentation here.

## Materials and Methods

### Library collection and sample annotation

The dataset comprised a diverse collection of human and *Drosophila* CUT&RUN libraries, three mixed CUT&RUN libraries generated for downsampling experiments, and four *Drosophila* Micro-C libraries. *Drosophila* H3K4me3, H3K27me3 and H3K36me3 CUT&RUN libraries in Fig. 3 were prepared from whole testes. The two human H3K4me3 CUT&RUN replicate-sampling series were generated from HEK293T and U2OS libraries, respectively. Of the four Micro-C libraries, Micro-C_1 and Micro-C_2 were prepared from S2 cells and Micro-C_3 and Micro-C_4 were from whole testes. Sources of all libraries are listed in Supplementary Table S1.

### CUT&RUN Library Preparation

CUT&RUN was performed using a published protocol(34) with modifications. For *Drosophila* testes, nuclei were released by Dounce homogenization in sucrose-containing buffer and collected by centrifugation. Nuclei were lightly fixed with 0.15% formaldehyde for 2 minutes and permeabilized with 0.1% Triton X-100 and used for subsequent steps after binding to Concanavalin A (ConA) coated beads. Antibodies against H3K4me3 (Millipore, cat. no. 07-473), H3K36me3 (Abcam, cat. no. ab9050) and H3K27me3 (Cell Signaling Technology, clone C36B11, cat. no. 9733) were used at a 1:100 dilution. Released DNA was purified and converted to sequencing libraries using NEBNext Ultra II DNA Library Prep Kit for Illumina (New England Biolabs, cat. no. E7645L), with final PCR amplification cycles ranging from 8 - 16.

### Micro-C Library Preparation

Micro-C libraries were prepared based on published Micro-C and Pico-C methods(29, 41), with modifications. Cells were fixed with 1% formaldehyde for 10 minutes, followed by 3 mM disuccinimidyl glutarate (DSG) for 45 minutes at room temperature. Isolated nuclei were immobilized on ConA-coated beads. MNase digestion conditions were optimized to yield approximately 90% mononucleosomes and 10% dinucleosomes, followed by end labeling with biotinylated nucleotides and proximity ligation. After reverse crosslinking, DNA was purified by phenol-chloroform extraction, enriched using streptavidin beads, and converted into sequencing libraries using NEBNext Ultra II DNA Library Prep Kit for Illumina (New England Biolabs, cat. no. E7645L). Fragments of approximately 400 bp were size-selected using solid-phase reversible immobilization (SPRI, Beckman Coulter, cat. no. A63881) beads to enrich for di-nucleosomal products.

### ONT Sequencing

Indexed libraries were quantified using the Qubit high-sensitivity assay (Thermo Fisher Scientific, cat. no. Q32854) and fragment size profiles were evaluated using the Agilent TapeStation High Sensitivity D5000 ScreenTape assay (Agilent Technologies, cat. no. 5067-5592). Purified PCR-amplified libraries were submitted to the Plasmidsaurus Premium PCR Sequencing service, using ONT native barcoding approach, which ligates adapters to the end of the molecules, preserving the full length sequences. Libraries were sequenced with ONT R10.4.1 flow cells.

### Library reconstruction

Each raw ONT read was scanned for inner TruSeq- or Nextera-compatible adapters. A reconstructed library unit was defined as a segment bounded by two inner adapters, and the sequence between the two adapters was defined as its insert. A single ONT read could contain more than one unit, and all units remained linked to their parent read. Inserts of ≤10 bp were classified as dimer-like, and longer inserts were retained. Outer P5/P7 sequences and index pairs were assessed separately from the inner adapters. Recovery and dimer-like fractions were calculated per raw read, and index-assignment fractions per reconstructed unit.

### Reconstruction of native electrophoresis

Because the exported files contained fluorescence measurements along the migration axis but did not provide a base-pair coordinate for each position, we converted migration position to fragment size using the TapeStation ladder. Eleven ladder peaks spanning 15-10,000 bp were identified, and the positions between these peaks were converted to fragment sizes by monotonic interpolation of log-transformed fragment length. Lower and upper reference marker peaks were used to confirm the calibration but were not used to adjust it.

Full-length Illumina library molecule lengths were reconstructed by adding the combined lengths of the sequencing adapters and indexes to the recovered insert lengths. Adapter lengths were determined based on the library preparation and indexing oligonucleotides used for each library (136 bp for libraries with the specified dual 8-bp indexing architecture). Two ONT-derived profiles were generated: an unweighted profile, in which each molecule contributed equally, and a length-weighted profile, in which each molecule contributed in proportion to its estimated total library length. Length weighting was used to approximate the DNA-mass-dependent fluorescence measured by TapeStation.

TapeStation and ONT-derived profiles were independently normalized to their maximum signal between 100 and 5,000 bp. Thus, these comparisons evaluate the shape of the fragment-size distributions rather than absolute DNA concentration or mass fraction. Gel-like representations were generated from the same profiles using 180 evenly spaced migration positions, with larger DNA fragments displayed toward the top of the lane. For quantitative comparison of profile shapes, distributions were area-normalized and Jensen–Shannon distance was calculated. Each ONT technical-repeat series was compared with its corresponding experimentally measured TapeStation profile.

The calibrated TapeStation traces were smoothed using a Gaussian kernel. The smoothing width was estimated from the widths of ladder peaks between 250 and 1,500 bp and corresponded to approximately 5.0-5.5 positions in the exported trace. The same calibration and smoothing parameters were used for all technical repeats within a given series.

### Construction and comparison of chromatin profiles

Reconstructed ONT library inserts were aligned to the reference genome using Bowtie2 with the --end-to-end and --very-sensitive settings(42). Only primary alignments to nuclear chromosomes with mapping quality scores (MAPQ) of ≥20 and <255 were retained. Alignments flagged as secondary, supplementary, failed quality control, or duplicates were excluded.

To compare chromatin enrichment profiles between ONT and Illumina sequencing, we generated normalized genomic coverage tracks and examined their signal distributions across defined genomic regions. ONT coverage was computed from reconstructed inserts and expressed as counts per million mapped reads (CPM) in 10-bp bins. For Illumina references, we used the 10-bp RPKM tracks produced by the Illumina processing pipeline, using the complete dataset and the original filtering, and converted them to CPM so that both platforms share the same units. For fixed-width bins, this conversion is a constant scaling (CPM = RPKM × bin length in kb). Profiles were generated with deepTools (v3.5.6) computeMatrix and plotProfile using 100-bp matrix bins(43). Pearson correlations between ONT and Illumina profiles were calculated over the same gene or region coordinates in both datasets. For gene-centered profiles, we used a common set of RefSeq gene coordinates for both sequencing platforms. Gene bodies were oriented according to transcriptional direction and scaled to a uniform length of 5 kb, with an additional 5 kb of unscaled genomic sequence upstream and downstream of each gene. Fixed gene sets contained 17,680 dm6 and 57,897 hg38 intervals, with missing values and genes shorter than one bin represented by zeros. For H3K27me3 profiles, we used a set of broad H3K27me3-enriched regions (4,606 broad peaks) identified from the corresponding full-depth Illumina sequencing dataset using MACS3 v3.0.4 in broad-peak mode (q-value cutoff, 0.05; broad-region cutoff, 0.1)(44). The same genomic regions were used to evaluate ONT and Illumina signals.

### Quantification of genomic signal concentration

To quantify the extent to which sequencing fragments were concentrated within specific genomic regions, we calculated a genomic concentration index based on the frequency of fragment pairs occurring within the same genomic bin. The analyzed genome was divided into nonoverlapping 10kb bins, and the number of fragment pair midpoints within each bin was counted. We then calculated the ratio of the observed number of fragment pairs occupying the same bin to the number expected if fragment midpoints were distributed uniformly across the analyzed genomic bases (collision observed/expected, O/E).

For N fragment midpoints, with *n_b_* in bin b, the observed number of same-bin pairs are:

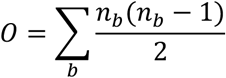

For an analyzed genome of length G and bins of width *w_b_*, the uniform collision probability and expected number of pairs are:

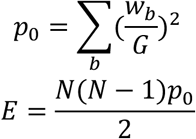

This metric is based on Simpson’s concentration index(45) and is equivalent to Morisita’s dispersion index for equal-width bins(27). An O/E value of 1 indicates the level of concentration expected under uniform random sampling, whereas values greater than 1 indicate increased genomic clustering of sequencing fragments.

### Comparison of Micro-C contact profiles and ligation-product structures

To compare Micro-C contact distributions between ONT and Illumina sequencing, terminal sequences from reconstructed ONT library inserts were aligned to *Drosophila melanogaster* genome (dm6) with BWA-MEM2(46, 47). For both platforms, contacts were retained only when both ends mapped to primary dm6 chromosomes with mapping quality (MAPQ) ≥30 and <255. ONT terminal alignments were additionally required to cover at least 80% of the query sequence. Contacts with identical genomic coordinates and strand orientations at both ends were considered duplicates and removed at 1-bp resolution. Inward-facing cis contacts separated by less than 200 bp were also excluded. Among the remaining unique, valid contacts, we calculated the fractions of cis contacts (both ends mapping to the same chromosome) and long-range cis contacts (cis contacts separated by ≥10 kb). Both fractions were calculated relative to all unique, valid contacts, including trans contacts (Table S3). These filtering criteria were used for cross-platform comparisons and were more stringent than the default MAPQ ≥20 threshold used in ONT-LibQC reports.

To examine the structure of individual Micro-C library molecules, we analyzed alignments across the entire reconstructed ONT insert to identify the genomic DNA fragments joined by ligation. Each insert was divided into consecutive genomic alignment blocks, with each block required to contain at least 40 aligned bases, MAPQ ≥20 and <255, and estimated sequence identity ≥85%. To be considered resolved, the identified blocks had to collectively cover at least 80% of the insert, with no more than 10 bp of overlap between blocks and no unexplained gaps longer than 30 bp. Adjacent blocks with collinear genomic alignments were merged. Inserts that did not meet these criteria were classified as unresolved. The proportions of molecules containing different numbers of blocks were calculated using all screened inserts, including unresolved molecules, but excluding adapter dimers. For molecules containing at least two resolved blocks, the length of each contributing genomic fragment was estimated from its aligned reference span. These block counts represent minimum estimates of the number of ligated fragments, because fragments mapping to adjacent genomic regions may be merged and short or ambiguously mapped fragments may not be detected. For Illumina libraries, genomic blocks were identified separately from each sequenced read end using the same alignment criteria. When both ends could be resolved, the larger block count was assigned to the read pair; otherwise, the pair was classified as unresolved. Thus, ONT block counts describe the complete reconstructed insert, whereas Illumina block counts reflect only the sequenced ends of each molecule.

To compare the performance of conventional paired-end Micro-C data processing, we generated artificial paired-end reads from ONT inserts by extracting up to 150 bp from each end, allowing the two sequences to overlap for inserts shorter than 300 bp. These ONT-derived read pairs and the original Illumina read pairs were processed using the same fastp, BWA-MEM2, and pairtools pipeline(31, 46, 48). The fractions of different pairtools mapping and processing categories were calculated relative to all parsed read pairs, both before and after duplicate marking (Extended Data Fig. 2B). To account for differences in sequencing depth, Illumina read pairs were randomly subsampled 50 times to match the number of parsed ONT read pairs, with duplicates identified separately within each subsample. ONT subsamples were analyzed using the same terminal-read pipeline, with duplicate marking repeated for each subsample (Extended Data Fig. 2C). For the full-depth reference datasets shown in Extended Data Fig. 2C, ONT and Illumina results were reported after and before duplicate marking, respectively.

### Reproducibility, computational and experimental downsampling analyses

To assess the reproducibility of ONT-LibQC with a limited number of sequencing reads, two human H3K4me3 CUT&RUN libraries (HEK293T and U2OS) were each sequenced in four separate ONT batches. Each ONT dataset was analyzed independently, and the resulting chromatin profiles were compared with those generated from the corresponding Illumina library.

To determine how sequencing depth affects the reproducibility of ONT-LibQC analysis, we performed computational downsampling of ONT reads. For each dataset, 50 random orderings of the raw reads were generated. From each ordering, nested subsets containing 250, 500, 1,000 and 2,000 reads were selected, such that each larger subset included all reads from the smaller ones. All reconstructed units and blocks from each selected read were retained, and CPM was recalculated for each subset. Subset profiles were generated by combining the coverage by the selected reads within deepTools genomic bins. The resulting downsampling profiles generated using bamCoverage and computeMatrix were compared to the original complete datasets. Chromatin profiles are presented as the mean ± standard deviation (SD) across 50 random samplings. Correlation coefficients and Micro-C metrics are summarized as medians with empirical 5th-95th percentile intervals unless stated otherwise (Supplementary Table 4). These intervals reflect sampling variability, not biological variation.

For experimental dilution, an H3K4me3 CUT&RUN library and an unrelated CUT&RUN library with different indices, both from U2OS cells, were mixed at volume ratios of 1:3, 2:2 and 3:1 (H3K4me3:unrelated). The i5 and i7 indexes were identified using the surrounding Illumina adapter sequences. Their combination was used to assign reconstructed inserts to source libraries, allowing for sequencing errors. H3K4me3 profiles were generated from reads whose reconstructed units were all assigned to the H3K4me3 index pair, yielding 506, 1,081 and 1,805 reads, respectively. Chromatin profiles were generated from these reads, with CPM normalization performed separately for each subset, and the resulting profiles were compared with the Illumina profile of the original, unmixed H3K4me3 library (Supplementary Table 4).

### Screening for potential spike-in sequences and mitochondrial fractions

To identify potential spike-in sequences, reconstructed inserts were aligned against a combined reference containing the host genome, candidate spike-in genomes, and sequencing controls (Extended Data Fig. 3A). *E. coli, S. cerevisiae, S. pombe* and *D. melanogaster* were included as candidate spike-in genomes, while lambda phage and PhiX were included as sequencing controls. Each insert was assigned to the reference with the best alignment if it met the following criteria: MAPQ ≥20 and <255, ≥80 aligned bases, ≥80% query coverage and ≥85% estimated sequence identity, and an alignment-score margin of ≥10 over the second-best alignment when one existed.

A candidate spike-in was considered detected when at least three qualifying inserts mapped to at least two distinct genomic regions, defined by the reference contig and 100-bp windows of alignment start positions. Candidates with fewer than three qualifying inserts were reported as traces if any were detected. Spike-ins originating from the same species as the host were marked as non-identifiable because their sequences could not be distinguished from the host genome. For each library, the number of qualifying inserts assigned to each candidate and the total number of screened inserts were recorded (Extended Data Fig. 3B). These signals were not used for normalization because no spike-in DNA was intentionally added to these libraries. Analyses were performed using Python, SAMtools(49) and Snakemake(21).

The mitochondrial fraction was the proportion of alignments to the mitochondrial genome among all alignments to the mitochondrial genome and the analyzed primary nuclear chromosomes. We counted only primary alignments with MAPQ ≥20 and <255, and excluded QC-fail and duplicate-flagged records. For CUT&RUN libraries, we used the alignments of reconstructed inserts; for Micro-C libraries, we used the alignments of the insert ends before contact selection. Each library or physical mixture is represented by its original ONT sequencing run (Extended Data Fig. 4).

## Use of AI tools

OpenAI Codex was used to assist with manuscript editing, code development and data analysis. All AI-assisted text, code and analyses were reviewed and verified by H.L. and J.J.K., who take full responsibility for the final content.

## Acknowledgements

We thank the Cornell University BRC Genomics Core Facility (RRID:SCR_021727) for next-generation sequencing services. We thank Boyuan Wang at University of Wisconsin-Madison, Jiacheng Wang at Duke University, Dr. Wei Sun at Virginia Tech and all members of Kim Lab for discussions and critical reading of the manuscript. We thank Dr. John T. Lis and Dr. Marcus Smolka at Cornell University, Dr. Margaret T. Fuller at Stanford University, FlyBase(50) and Bloomington Drosophila Stock Center (BDSC; NIH P40OD018537) for materials and resources.

## Data and code availability

Relevant numerical data are provided in the Supplementary Information. Sequencing data supporting this study have been deposited in the Gene Expression Omnibus under accession number GSE350270. Source code is available at https://github.com/Shall-We-Dance/ONT-LibQC, and the version associated with this publication is archived on Zenodo (DOI: 10.5281/zenodo.23481973). The web portal is available at https://ontlibqc.org.

## Author contributions

J.J.K. and H.L. conceived the study. H.L. designed and performed the experiments, developed the analysis pipeline, and established and maintained the web portal. J.J.K. supervised and supported the project. Both authors wrote and revised the manuscript.

---

**Extended Data Figure 1.**
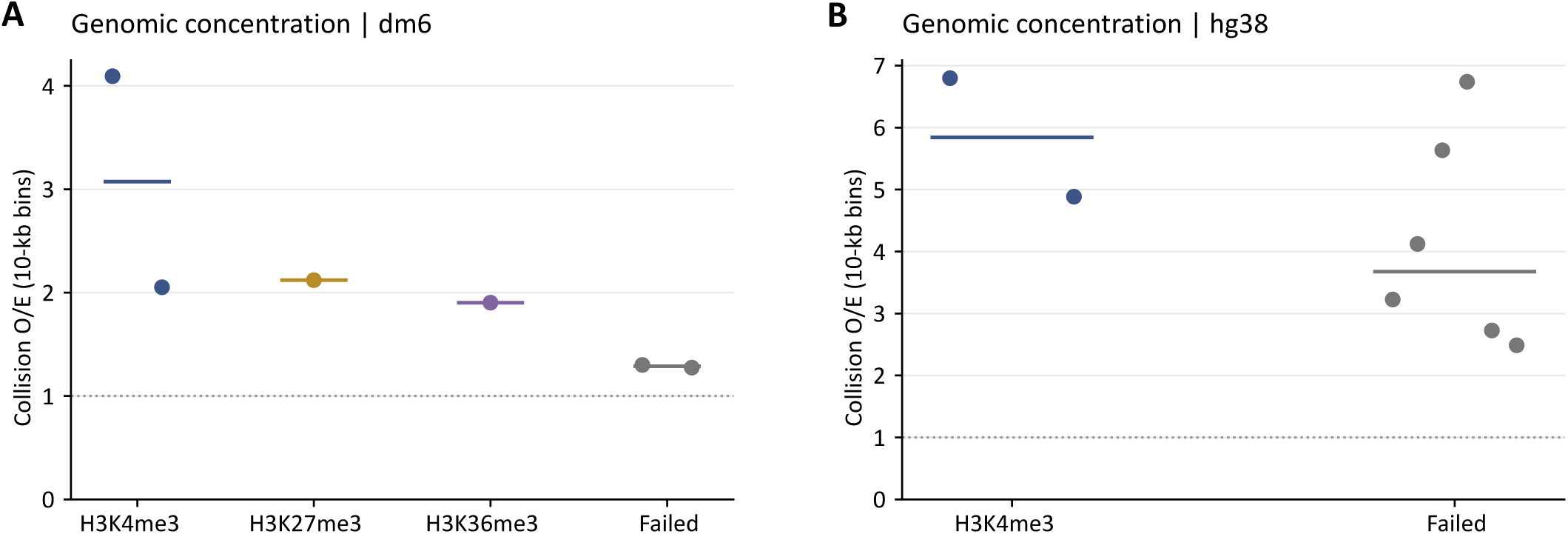
Genomic concentration across chromatin libraries. **A,B,** Collision O/E in 10-kb bins for *Drosophila* (dm6) and human (hg38) libraries. Points represent individual libraries, horizontal bars indicate group medians, and dotted lines represent uniform genomic occupancy.

**Extended Data Figure 2.**
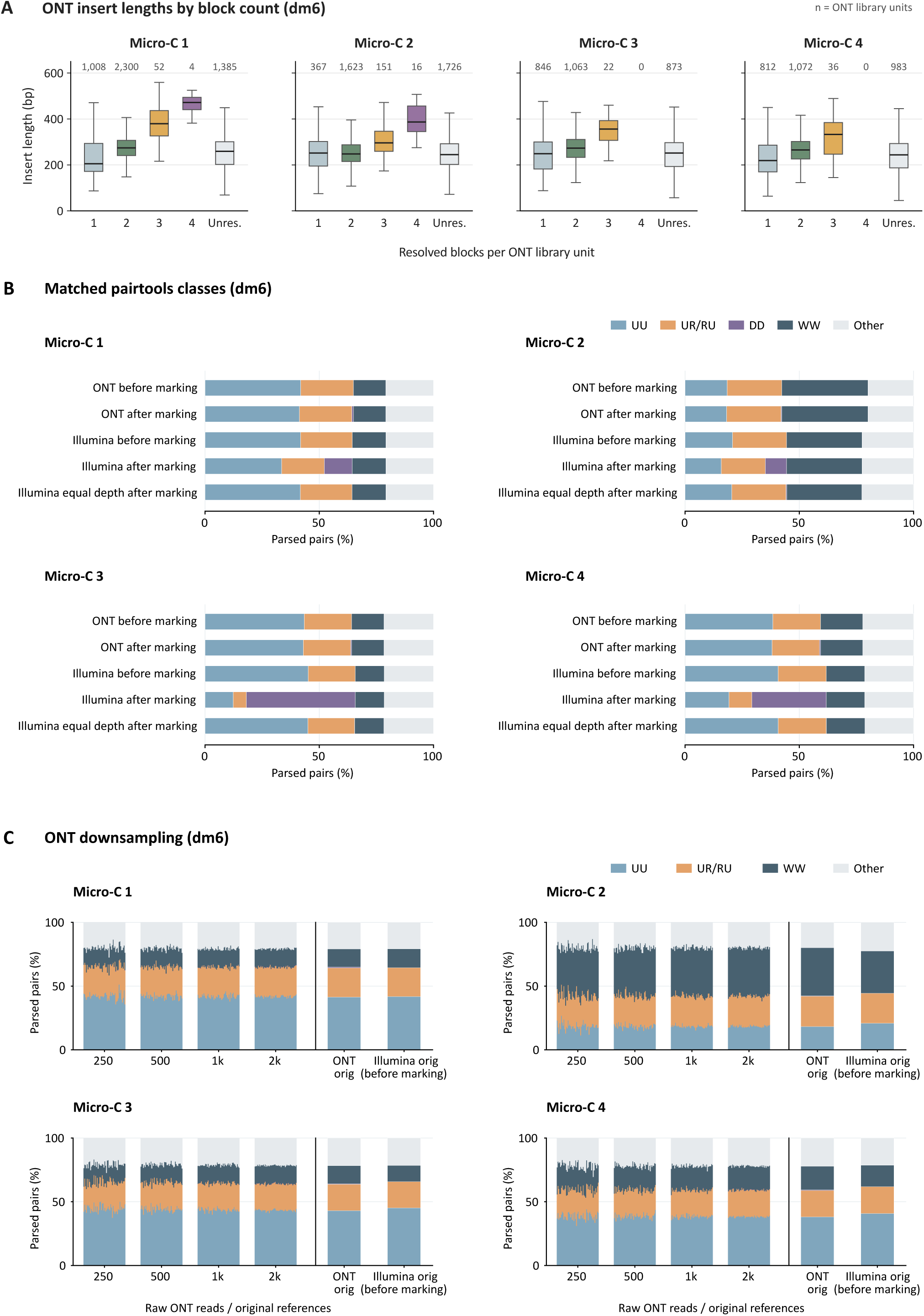
Micro-C read-pair classes under duplicate marking and downsampling, and insert lengths by resolved block count. **A,** Insert lengths of ONT library units in the four Micro-C libraries, grouped by the number of resolved genomic blocks. Adapter dimers are excluded. Boxes show the median and interquartile range, and whiskers extend to 1.5 times the interquartile range. Numbers above the boxes are unit counts. Unres., unresolved units. **B,** ONT and Illumina read-pair classifications before and after duplicate marking, along with Illumina data downsampled to match the number of parsed ONT pairs. Depth-matched Illumina bars show the mean of 50 random draws, with duplicates marked separately within each draw. **C,** Thin bars represent 50 nested ONT subsets at each sequencing depth; wider bars represent the original, full-depth ONT and Illumina datasets. In **B** and **C**, fractions were calculated using all parsed pairs as the denominator. UU, uniquely mapped pairs; UR/RU, rescued pairs; DD, coordinate duplicates; WW, masked complex walks; Other, remaining outcomes.

**Extended Data Figure 3.**
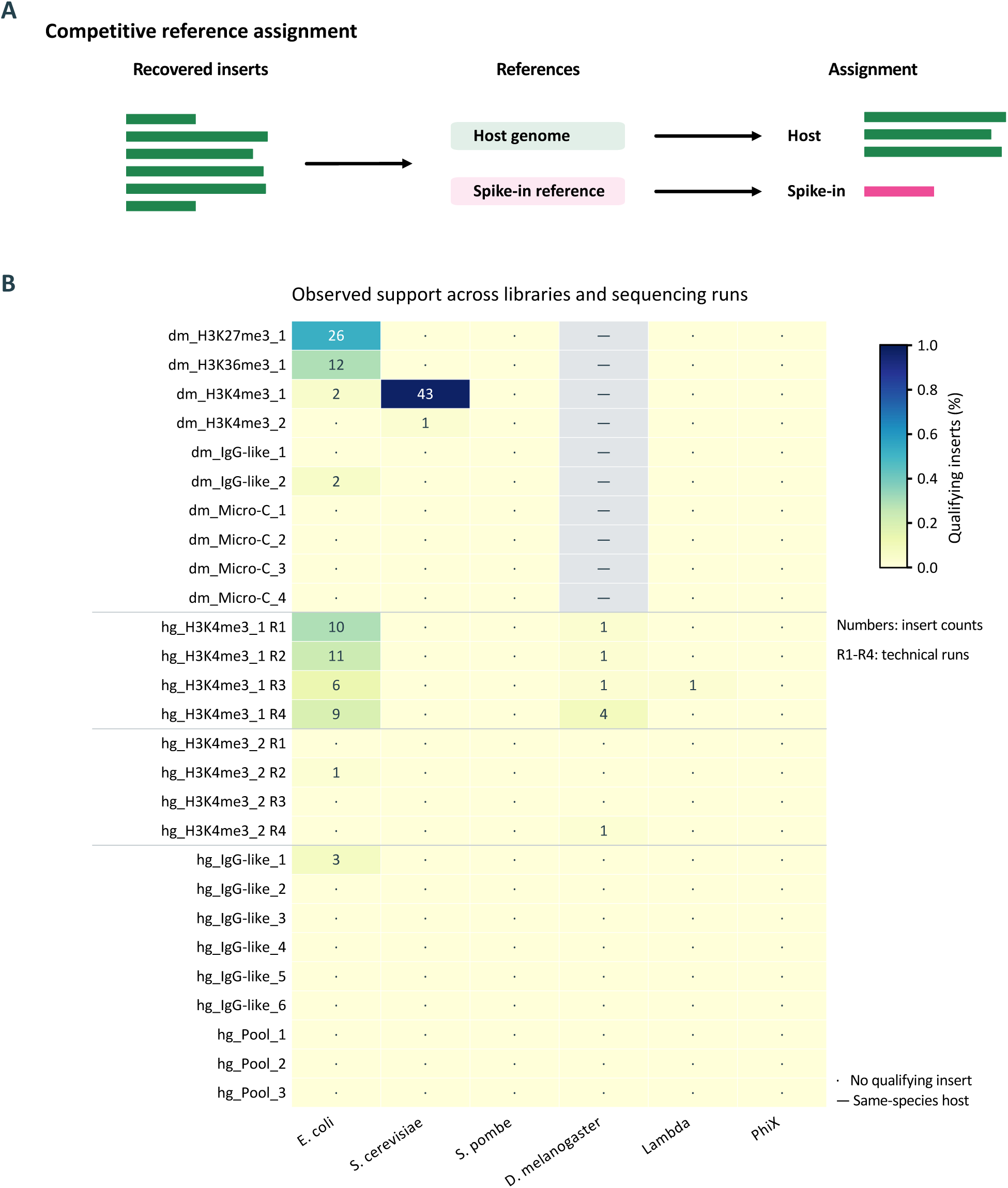
Screening for candidate spike-in sequences. **A,** Reconstructed inserts were screened using a combined reference containing host genome and candidate spike-in sequences. **B,** Percentages of qualifying inserts across the 21 original datasets in 27 sequencing runs. Labels indicate read counts, dots represent zero counts, and gray cells indicate candidates that cannot be distinguished from the host genome. Lambda and PhiX were sequencing controls. No spike-in DNA was intentionally added to these libraries, and these results were not used for normalization. Counts and denominators are provided in Table S5.

**Extended Data Fig. 4.**
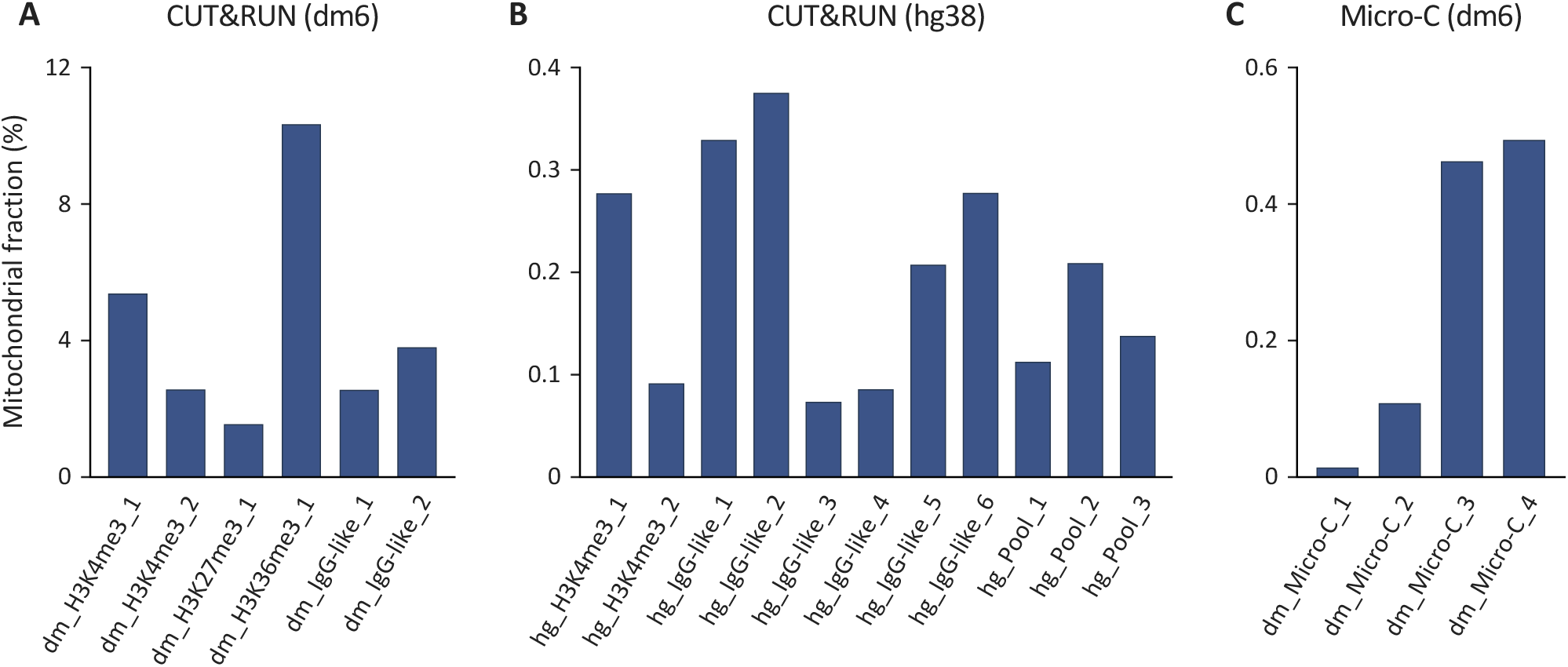
Mitochondrial fractions across ONT libraries. **A,B,** Mitochondrial fractions in (A) *Drosophila* (dm6) and (B) human (hg38) CUT&RUN libraries. hg_Pool_1-3 are the three physical mixtures. **C,** Mitochondrial fractions in four *Drosophila* Micro-C libraries. Each fraction is the number of mitochondrial alignments divided by the number of mitochondrial plus primary nuclear alignments passing the mapping filters (Methods). CUT&RUN fractions are based on alignments of reconstructed inserts, and Micro-C fractions on alignments of the insert ends before contact selection. Each bar shows the original ONT sequencing run of one library or pool. Y-axis scales differ between panels.

## References

1. Hainer, S.J., Bošković, A., McCannell, K.N., et al. (2019) Profiling of Pluripotency Factors in Single Cells and Early Embryos. Cell, 177, 1319–1329.e11.

2. Kaya-Okur, H.S., Wu, S.J., Codomo, C.A., et al. (2019) CUT&Tag for efficient epigenomic profiling of small samples and single cells. Nat Commun, 10, 1930.

3. Brind’Amour, J., Liu, S., Hudson, M., et al. (2015) An ultra-low-input native ChIP-seq protocol for genome-wide profiling of rare cell populations. Nat Commun, 6, 6033.

4. Emerson, F.J. and Lee, S.S. (2022) CUT&RUN for Chromatin Profiling in Caenorhabditis elegans. Curr Protoc, 2, e445.

5. Bartosovic, M., Kabbe, M. and Castelo-Branco, G. (2021) Single-cell CUT&Tag profiles histone modifications and transcription factors in complex tissues. Nat Biotechnol, 39, 825–835.

6. Cejas, P., Li, L., O’Neill, N.K., et al. (2016) Chromatin immunoprecipitation from fixed clinical tissues reveals tumor-specific enhancer profiles. Nat Med, 22, 685–691.

7. Zhang, H., Polavarapu, V.K., Xing, P., et al. (2022) Profiling chromatin accessibility in formalin-fixed paraffin-embedded samples. Genome Res, 32, 150–161.

8. Chu, T., Rice, E.J., Booth, G.T., et al. (2018) Chromatin run-on and sequencing maps the transcriptional regulatory landscape of glioblastoma multiforme. Nat Genet, 50, 1553–1564.

9. Corces, M.R., Trevino, A.E., Hamilton, E.G., et al. (2017) An improved ATAC-seq protocol reduces background and enables interrogation of frozen tissues. Nat Methods, 14, 959–962.

10. Hayashi, R. and Inoue, A. (2023) Low-Input CUT&RUN for Mouse Oocytes and Preimplantation Embryos. Methods Mol Biol, 2577, 83–92.

11. Ruiz, S.E., Maul, R.W. and Gearhart, P.J. (2025) Optimized CUT&RUN protocol for activated primary mouse B cells. PLoS One, 20, e0322139.

12. Meers, M.P., Bryson, T.D., Henikoff, J.G., et al. (2019) Improved CUT&RUN chromatin profiling tools. Elife, 8, e46314.

13. Goel, V.Y., Huseyin, M.K. and Hansen, A.S. (2023) Region Capture Micro-C reveals coalescence of enhancers and promoters into nested microcompartments. Nat Genet, 55, 1048–1056.

14. Ou, J., Liu, H., Yu, J., et al. (2018) ATACseqQC: a Bioconductor package for post-alignment quality assessment of ATAC-seq data. BMC Genomics, 19, 169.

15. Grandi, F.C., Modi, H., Kampman, L., et al. (2022) Chromatin accessibility profiling by ATAC-seq. Nat Protoc, 17, 1518–1552.

16. Haring, M., Offermann, S., Danker, T., et al. (2007) Chromatin immunoprecipitation: optimization, quantitative analysis and data normalization. Plant Methods, 3, 11.

17. Zee, A., Deng, D.Z.Q., Adams, M., et al. (2022) Sequencing Illumina libraries at high accuracy on the ONT MinION using R2C2. Genome Res, 32, 2092–2106.

18. Mumm, C., Drexel, M.L., McDonald, T.L., et al. (2023) Multiplexed long-read plasmid validation and analysis using OnRamp. Genome Res, 33, 741–749.

19. Li, W., Miller, D., Liu, X., et al. (2024) Arrayed in vivo barcoding for multiplexed sequence verification of plasmid DNA and demultiplexing of pooled libraries. Nucleic Acids Res, 52, e47.

20. Oxford Nanopore Technologies (2026) Ligation sequencing amplicons - Native Barcoding Kit 24 V14 (SQK-NBD114.24). Oxford Nanopore Technologies.

21. Mölder, F., Jablonski, K.P., Letcher, B., et al. (2021) Sustainable data analysis with Snakemake. F1000Res, 10, 33.

22. Quail, M.A., Swerdlow, H. and Turner, D.J. (2009) Improved protocols for the illumina genome analyzer sequencing system. Curr Protoc Hum Genet, Chapter 18, Unit 18.2.

23. Knapp, M., Stiller, M. and Meyer, M. (2012) Generating barcoded libraries for multiplex high-throughput sequencing. Methods Mol Biol, 840, 155–170.

24. Thompson, J.R., Marcelino, L.A. and Polz, M.F. (2002) Heteroduplexes in mixed-template amplifications: formation, consequence and elimination by ‘reconditioning PCR’. Nucleic Acids Res, 30, 2083–2088.

25. Barski, A., Cuddapah, S., Cui, K., et al. (2007) High-resolution profiling of histone methylations in the human genome. Cell, 129, 823–837.

26. Kharchenko, P.V., Alekseyenko, A.A., Schwartz, Y.B., et al. (2011) Comprehensive analysis of the chromatin landscape in Drosophila melanogaster. Nature, 471, 480–485.

27. Morisita, M. (1962) *I*_σ -_Index, a measure of dispersion of individuals. Population Ecology, 4, 1–7.

28. Lieberman-Aiden, E., van Berkum, N.L., Williams, L., et al. (2009) Comprehensive mapping of long-range interactions reveals folding principles of the human genome. Science, 326, 289–293.

29. Hsieh, T.-H.S., Weiner, A., Lajoie, B., et al. (2015) Mapping Nucleosome Resolution Chromosome Folding in Yeast by Micro-C. Cell, 162, 108–119.

30. Deshpande, A.S., Ulahannan, N., Pendleton, M., et al. (2022) Identifying synergistic high-order 3D chromatin conformations from genome-scale nanopore concatemer sequencing. Nat Biotechnol, 40, 1488–1499.

31. Open2C, Abdennur, N., Fudenberg, G., et al. (2024) Pairtools: From sequencing data to chromosome contacts. PLoS Comput Biol, 20, e1012164.

32. Daley, T. and Smith, A.D. (2013) Predicting the molecular complexity of sequencing libraries. Nat Methods, 10, 325–327.

33. DeMaere, M.Z. and Darling, A.E. (2021) qc3C: Reference-free quality control for Hi-C sequencing data. PLoS Comput Biol, 17, e1008839.

34. Skene, P.J. and Henikoff, S. (2017) An efficient targeted nuclease strategy for high-resolution mapping of DNA binding sites. eLife, 6, e21856.

35. Landt, S.G., Marinov, G.K., Kundaje, A., et al. (2012) ChIP-seq guidelines and practices of the ENCODE and modENCODE consortia. Genome Res., 22, 1813–1831.

36. Fursova, N.A., Turberfield, A.H., Blackledge, N.P., et al. (2021) BAP1 constrains pervasive H2AK119ub1 to control the transcriptional potential of the genome. Genes Dev., 35, 749–770.

37. Ferrari, K.J., Scelfo, A., Jammula, S., et al. (2014) Polycomb-Dependent H3K27me1 and H3K27me2 Regulate Active Transcription and Enhancer Fidelity. Molecular Cell, 53, 49–62.

38. Daley, T. and Smith, A.D. (2014) Modeling genome coverage in single-cell sequencing. Bioinformatics, 30, 3159–3165.

39. Patel, L.A., Cao, Y., Mendenhall, E.M., et al. (2024) The Wild West of spike-in normalization. Nat Biotechnol, 42, 1343–1349.

40. Aird, D., Ross, M.G., Chen, W.-S., et al. (2011) Analyzing and minimizing PCR amplification bias in Illumina sequencing libraries. Genome Biol, 12, R18.

41. Maziak, N., Zhang, Y., Groll, F., et al. (2026) Three-dimensional genome reorganization foreshadows zygotic genome activation in Drosophila. Nat Genet, 58, 607–617.

42. Langmead, B. and Salzberg, S.L. (2012) Fast gapped-read alignment with Bowtie 2. Nat Methods, 9, 357–359.

43. Ramírez, F., Ryan, D.P., Grüning, B., et al. (2016) deepTools2: a next generation web server for deep-sequencing data analysis. Nucleic Acids Res, 44, W160–W165.

44. Doherty, P., Hu, Q., Zhuang, Z., et al. (2026) MACS3: A Peak-calling Platform for Bulk and Single-cell Regulatory Genomics. Genomics, Proteomics & Bioinformatics, 10.1093/gpbjnl/qzag097.

45. Simpson, E.H. (1949) Measurement of Diversity. Nature, 163, 688–688.

46. Vasimuddin, Md., Misra, S., Li, H., et al. (2019) Efficient Architecture-Aware Acceleration of BWA-MEM for Multicore Systems. In 2019 IEEE International Parallel and Distributed Processing Symposium (IPDPS). IEEE, Rio de Janeiro, Brazil, pp. 314–324.

47. Li, H. and Durbin, R. (2009) Fast and accurate short read alignment with Burrows–Wheeler transform. Bioinformatics, 25, 1754–1760.

48. Chen, S., Zhou, Y., Chen, Y., et al. (2018) fastp: an ultra-fast all-in-one FASTQ preprocessor. Bioinformatics, 34, i884–i890.

49. Danecek, P., Bonfield, J.K., Liddle, J., et al. (2021) Twelve years of SAMtools and BCFtools. GigaScience, 10, giab008.

50. Öztürk-Çolak, A., Marygold, S.J., Antonazzo, G., et al. (2024) FlyBase: updates to the Drosophila genes and genomes database. Genetics, 227, iyad211.

